# Behavioural analysis of single-cell aneural ciliate, *Stentor roeselii*, using machine learning approaches

**DOI:** 10.1101/594796

**Authors:** Kiều Mi Trịnh, Matthew T. Wayland, Sudhakaran Prabakaran

**Affiliations:** Trinity College, University of Cambridge, CB2 1TQ, UK; Department of Genetics, University of Cambridge, Downing Site, CB2 3EH, UK; Department of Zoology, University of Cambridge, Downing Street, Cambridge, CB2 3EJ, UK; Department of Biology, Indian Institute of Science Education and Research, Pune, Maharashtra, 411008, India; St Edmund’s College, University of Cambridge, CB3 0BN, UK

## Abstract

There is still a significant gap between our understanding of neural circuits and the behaviours they compute – i.e. the computations performed by these neural networks (Carandini 2012). Learning, behaviour, and memory formation, what used to only be associated with animals with neural systems, have been observed in many unicellular aneural species, namely Physarum, Paramecium, and Stentor (Tang & Marshall 2018). As these are fully functioning organisms, yet being unicellular, there is a much better chance to elucidate the detailed mechanisms underlying these learning processes in these organisms without the complications of highly interconnected neural circuits. An intriguing learning behaviour observed in *Stentor roeselii* (Jennings 1902) when stimulated with carmine has left scientists puzzled for more than a century. So far, none of the existing learning paradigm can fully encapsulate this particular series of five characteristic avoidant reactions. Although we were able to observe all responses described in literature and in a previous study (Dexter et al. 2019, manuscript in preparation), they do not conform to any particular learning model. We then investigated whether models based on machine learning approaches, including decision tree, random forest, and feed-forward neural networks could infer and predict the behavior of *S. roeselii*. Our results showed that an artificial neural network with multiple ‘computational’ neurons is inefficient at modelling the single-celled ciliate’s avoidant reactions. This has highlighted the complexity of behaviours in aneural organisms. Additionally, this report will also discuss the significance of elucidating molecular details underlying learning and decision-making processes in these unicellular organisms, which could offer valuable insights that are applicable to higher animals.

---

Since the 1700s many behaviours observed in lower unicellular organisms, such as *Physarum, Paramecium*, or *Stentor*, have been successfully demonstrated to satisfy many of the existing learning paradigms, from simple non-associative (Boisseau et al. 2016; Osborn et al. 1973; Tang & Marshall 2018; Wood 1973; Eisenstein 1975) to more complex associative models (Hennessey et al. 1979; Shirakawa et al. 2011). These observations have left scientists puzzled. To what extent do these organisms possess an awareness of their surrounding? Is it at all comparable to that experienced by higher animals? Herbert Jennings – one of the most influential biologists in the field of behaviours in aneural organisms – published some very detailed written accounts of the unique response seen in the single-cell ciliate *Stentor roeselii* upon mechanical or chemical stimulation (Jennings 1902). In particular, when stimulated with carmine particles (Fig. 1), the organisms were described to perform a series of five characteristic observable avoidant reactions in order to remove themselves (or ‘to leave’) from the noxious stimulus, provided that the particles were persistently present in the surrounding environment. This complicated sequence of action was acknowledged as one of “the most intricate behaviors so far recorded in unicellular animals” by Dennis Bray (Bray 2009). The five reactions are generally seen to occur in a particular order (Fig. 2), albeit with several variations. Sometimes the typical response order is not strictly followed, or time taken to switch between different avoidant reactions varies widely from one organism to another. Jennings was able to demonstrate that these differences in response are not due to fatigue, and thus concluded that the organism had performed some form of complex learning; an altered response due to prior experience (Jennings 1902).

**Fig 1.**
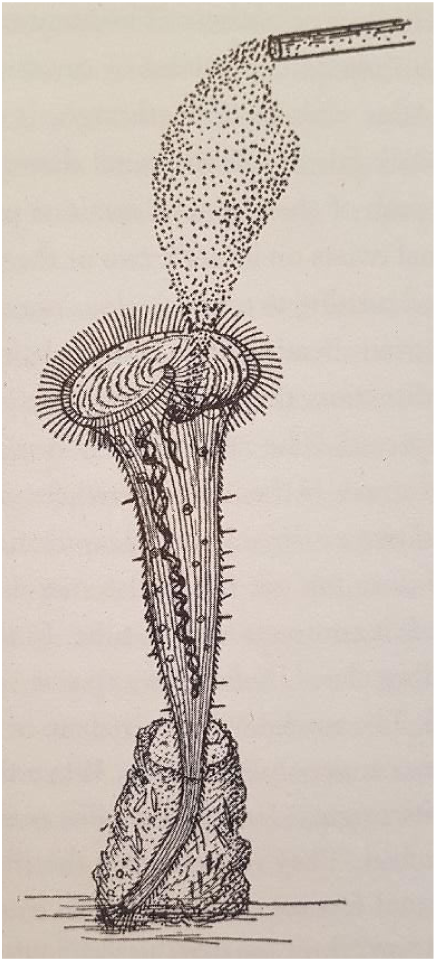
A sketch of carmine particles introduced to the buccal cavity of a *S. roeselii*. Illustration of the experiment from Jennings’ paper (Jennings 1906). Carmine particles are released over the mouth of a *S. roeselii* that are attached to a surface via its tube and holdfast

**Fig 2.**
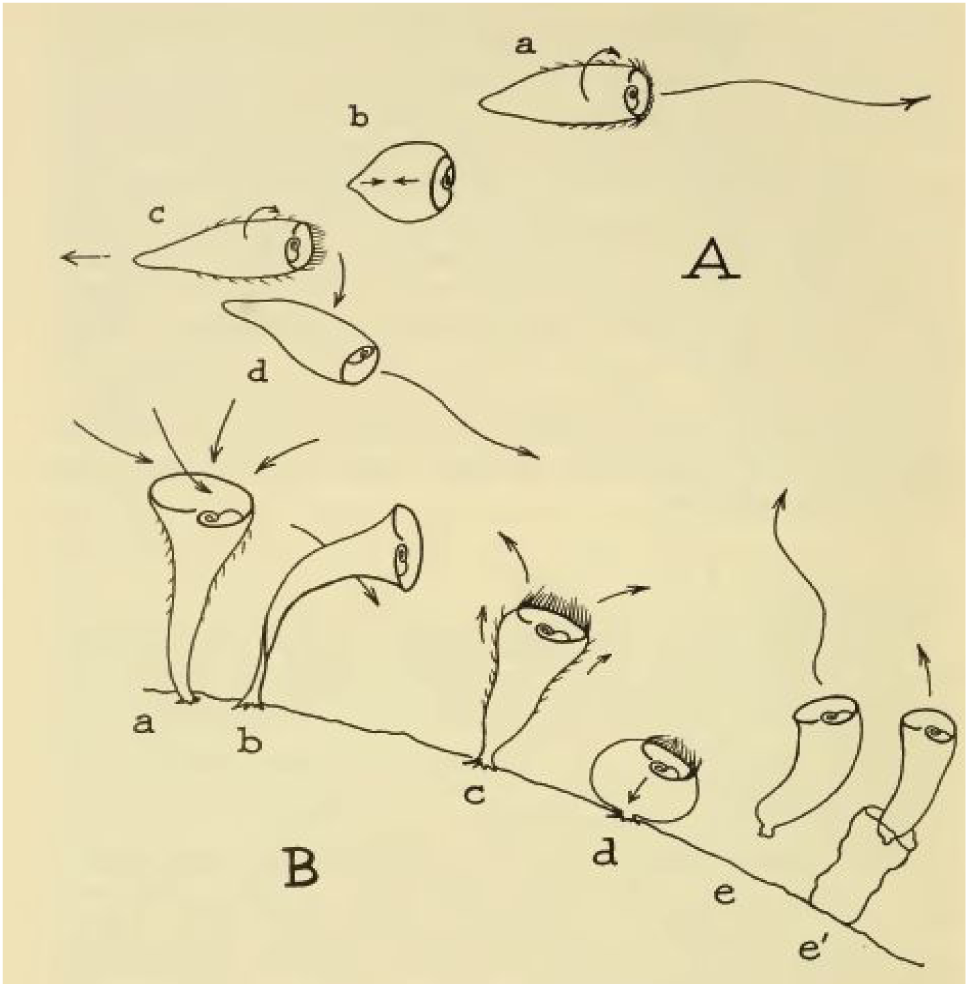
Series of five avoidant reactions observed in *S. roeselii*. This is Vance Tartar’s illustration of the five reactions of *S. roeselii* stimulated by carmine in the order described by Jennings (Tartar 1961). As long as the stimulus is still present in the surrounding environment, these five reactions are: a. No response – at rest b. Bending away from the source c. Transient stop of cilia beating, reversal of spiraling direction d. Strong full contraction e. Detachment and swim away

Multiple attempts to characterize this intriguing behaviour were later carried out, some of which challenged Jennings’ proposition (Reynierse & Walsh 1967; Wood 1969). However, some of these subsequent experiments used a related but more motile species – *S. coeruleus*. Thus, the exact Jennings observations were not seen and hence the avoidance behaviours were considered irreproducible. In an earlier (Dexter et al. 2019, manuscript in preparation), we developed a video-microscopy experimental paradigm and used *S. roeselii* to duplicate Jennings’ experiments. This study verified Jennings’ findings of a complex hierarchy of avoidance behaviours, which indicated a complex decision making process underlying the behaviours.

All these earlier studies were able to demonstrate that the observations in *S. roeselii* did not conform to any existing learning model for single cell organisms - they did not indicate habituation nor adaptive sensitization (which may require more than a life time to acquire!). Staddon had earlier suggested that it could be operant behaviour – behaviour “guided by its consequences”, and proposed some possible mechanisms, yet, no concrete conclusion was made (Staddon 1983).

Since there is no current learning model that can fully explain the series of avoidant reactions in *S. roeselii*, we examined if decision ‘to leave’ could be predicted from time spent performing each of the avoidant reactions using models based on Decision Tree, Random Forest, and Artificial Neural Network (ANN) machine learning algorithms. ANNs have proven their power with notable successes in applications across numerous fields, including modeling complex cognitive activities in higher animals (Savelli & Knierim 2018; Yang et al. 2019). We set out to explore how effective ANNs are at predicting *S. roeselii*’s behaviour, and are particularly interested to find out the number of computational neurons required for an ANN to be proficient at predicting the behaviour observed in a single-celled organism.

## Replication of Jennings’ original experiment – heterogeneity in S. roeselii’s behaviour

We replicated Jenning’s experiment and validated the complexity of the observed behaviours by (Dexter et al. 2019, manuscript in preparation). Upon stimulation with red-fluorescent latex beads, we were able to observe all the five avoidant reactions described by Jennings. Fig. S1 shows our simple experimental set-up. We used a light microscope to observe and record *S. roeselii’s* behaviour and a gravity based water reservoir was used for pulsed bead stimulations. Since observations provided by Jennings were only qualitative description via words and sketches, with experimental methods not being documented in detail, all observations were subjected to our own interpretation as illustrated in Fig. 3. The high level of heterogeneity in *S. roeselii*’s behaviour upon stimulation noted by Jennings (Jennings 1902) were also detected. The order of the series of reaction is not always the same as that illustrated in Fig. 2. In many instances, halting and reversing in direction of cilia beating took place before obvious bending was seen. The extent to which bending movements were performed varied massively. Generally, each of the five responses was repeated for a while before the organism decided to move on to the next, with large differences in the number of repeats for each response between organisms. Moreover, many individual *S. roeselii* did not demonstrate all five reactions, with some omitting bending, some contraction, and others immediately detaching. Occasionally, some *S. roeselii* immediately contracted upon stimulation. However, this could have been a reaction to the strong water pressure from releasing the beads into the environment.

**Fig 3.**
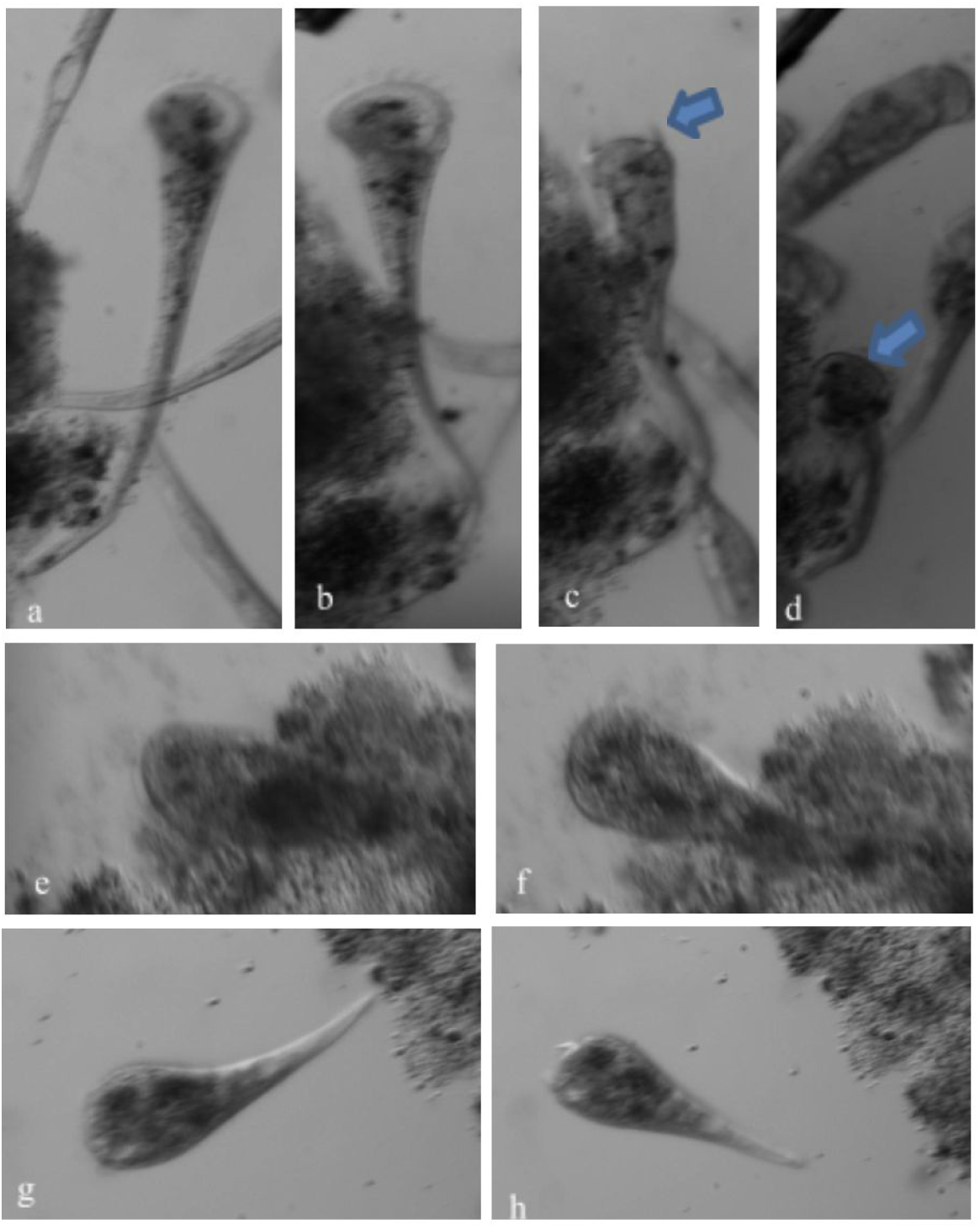
Example observations of *S. roeselii*’s different behavioural responses upon stimulation with polystyrene beads. At rest: *S.roeselii* is fully extended with cilia beating to generate a vortex current a. Bending b. Transient halting of cilia beating (arrow) c. Full contraction after encountering the fluorescent beads e-f. The organism slowly extended after contraction until full length is reached g-h. Detachment

Performing the experiment repeatedly with absolute consistency of control variables was challenging. This may or may not have affected the *Stentor*s’ behavior across experiments. For instance, experiments typically involved recording over five sessile *S. roeselii* simultaneously due to time limitations. Thus, it can be hard to distinguish if contraction of these organisms was a response to stimulation or due to collision with others swimming in close vicinity. Additionally, the amount of beads released into the environment for each experiment was, though roughly the same, not exact.

Analysis of the video recordings was not always straightforward. Cilia movement was only clearly observable when the organisms were positioned correctly in the plane of focus. Quite often, the organism bended in multiple directions throughout the course of the experiment, and therefore, their cilia were not always observable. Moreover, the reversal of cilia beating is usually accompanied by a twist or slight bending, which could lead to potential mis-classification of response if cilia were not observable. Another factor that could impact the quality of input variables is that the videos collected are of different length (see Material Methods). In an attempt to mitigate these factors, experiments where these issues were more pronounced were discarded from analysis. Being aware of all complications and the wide variation in *S. roeselii*’s behaviours, we were extremely cautious and very careful to make the most consistent and quantitative analysis possible.

## Evidence against Operant behaviour

Operant behaviour is described as a form of goal-directed behaviour. This learning model suggests that *S. roeselii*’s decision of switching from one response to the next is underpinned by a mechanism that “compute[s] the relative importance of time [spent repeating a particular response] and concentration [of the noxious substance in its vicinity]” (Staddon 1983). Staddon suggested a mechanism whereby the five reactions were graded according to their cost (energy expenditure +/-forgoing the opportunity to obtain further food), with each successive reaction only occurring above a certain threshold concentration of noxious beads. Additionally, each threshold elevates as the reaction continues to occur – ie. a form of habituation – and eventually, will overtake that of the next reaction in the series, resulting to a switch in behaviour. However, this seems to imply that the organism needs to go through the whole sequence of five reactions in the exact order every time it is stimulated. Yet, this was not always in our experiments. There were multiple instances where the first two or three stages were skipped, or the order of reactions was not followed. Staddon also mentioned that the above mechanism was just one of the many possibilities one could come up with. Ultimately, he conceded that the exact operant behaviour mechanism could not be verified without further physiological and behavioral analysis.

## Modelling S. roeselii’s avoidant behaviour using machine learning approaches

We decided to use the time (in seconds) spent performing each of the avoidant reactions described by Jennings as features to train our machine learning models. This includes the duration of (1) being at rest, (2) bending, (3) cilia reversal and (4) contraction (Video 1). Detachment was used as an outcome in our model, and so the 5^th^ feature included was (5) number of contractions observed. There are many more features which can be extracted from the raw data collected from analyzing the videos, such as dynamics of contraction and retraction, time taken for each contraction, or order of events taking place, etc. Nonetheless, as the data set is relatively small, it is not appropriate to use too many features to train the models.

The correlations between these features were investigated (Fig. 4). The results showed that contraction time and number of contraction is highly positively correlated, which is to be expected as the longer the organism spent in contraction stage, the more opportunities they would have to perform contraction. With respect to the outcome “Leave”, the duration of contraction and cilia reversal stages are the most negatively correlated features, with the number of contractions having a slightly less negative correlation.

**Fig 4.**
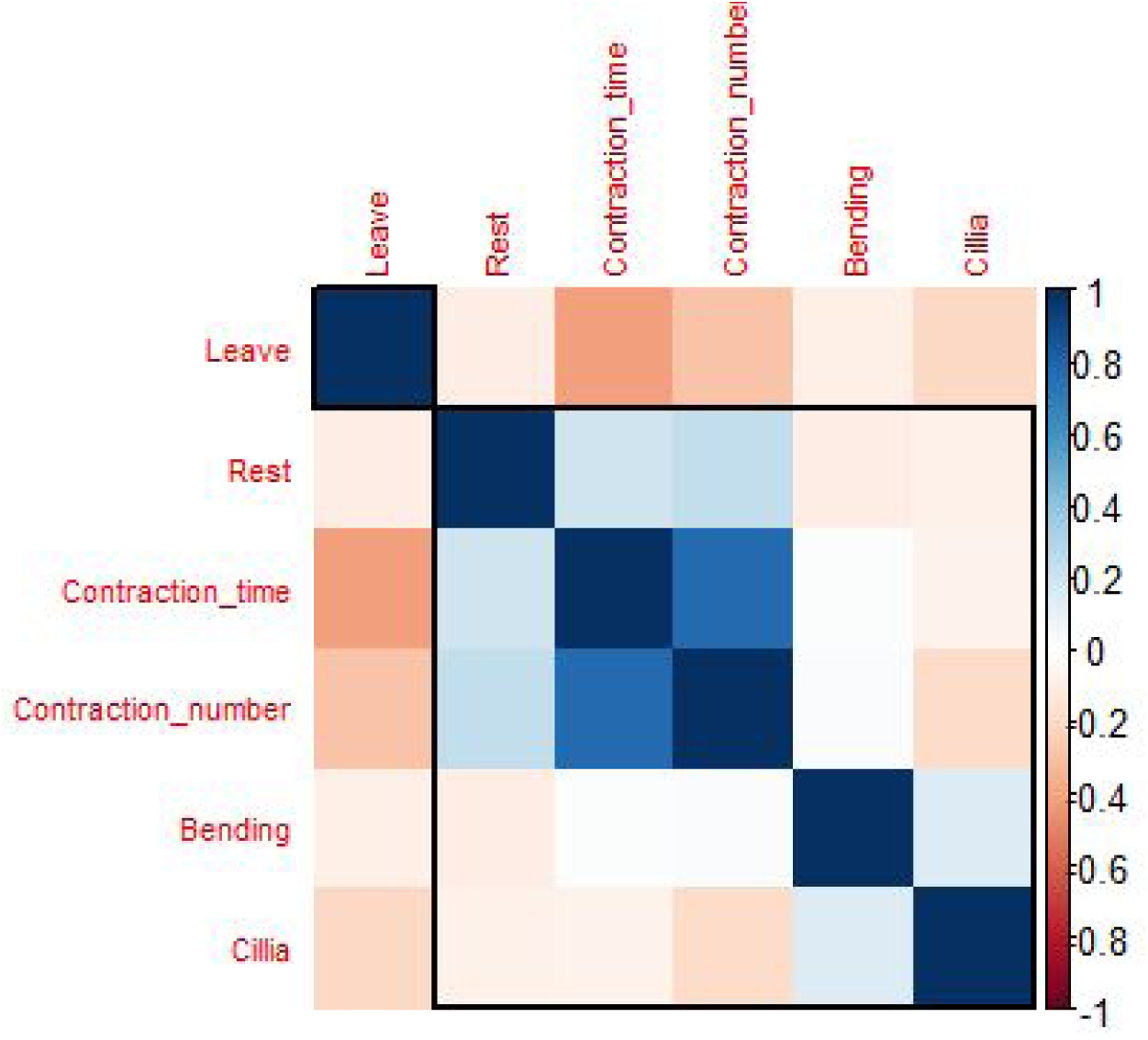
Correlation between 5 input features and output. The five input features that were used in the analysis are time spent at rest (Rest), bending (Bending), halting and reversing cilia (Cilia), contraction (Contraction_time) and number of contraction (Contraction_number) and output is determined as whether or not the organism detached and swam away (Leave). The correlation plot shows Contraction_time, Contraction_number, and Cilia being negatively correlated (orange/ light pink) to the outcome (in order of decreasing correlation strength – indicated by the decrease in the colour darkness)

The first classification model was based on Decision Tree algorithm. The tree-like flowchart (Fig. 5) was generated, with each internal node representing a “test” on an attribute, and the outcome of the test – the decision – is displayed on the branch. The decision is made at each branch until it comes to the termination point. At each node, data points are initially segregated based on all input variables individually, and the split that generates the most homogeneous classes will be chosen and displayed. Thus, the decision tree model identifies the significance hierarchy of input features used by the model after being trained. Applying this algorithm on our data set, contraction time and cilia reversal time were recognized as the most critical features.

**Fig 5.**
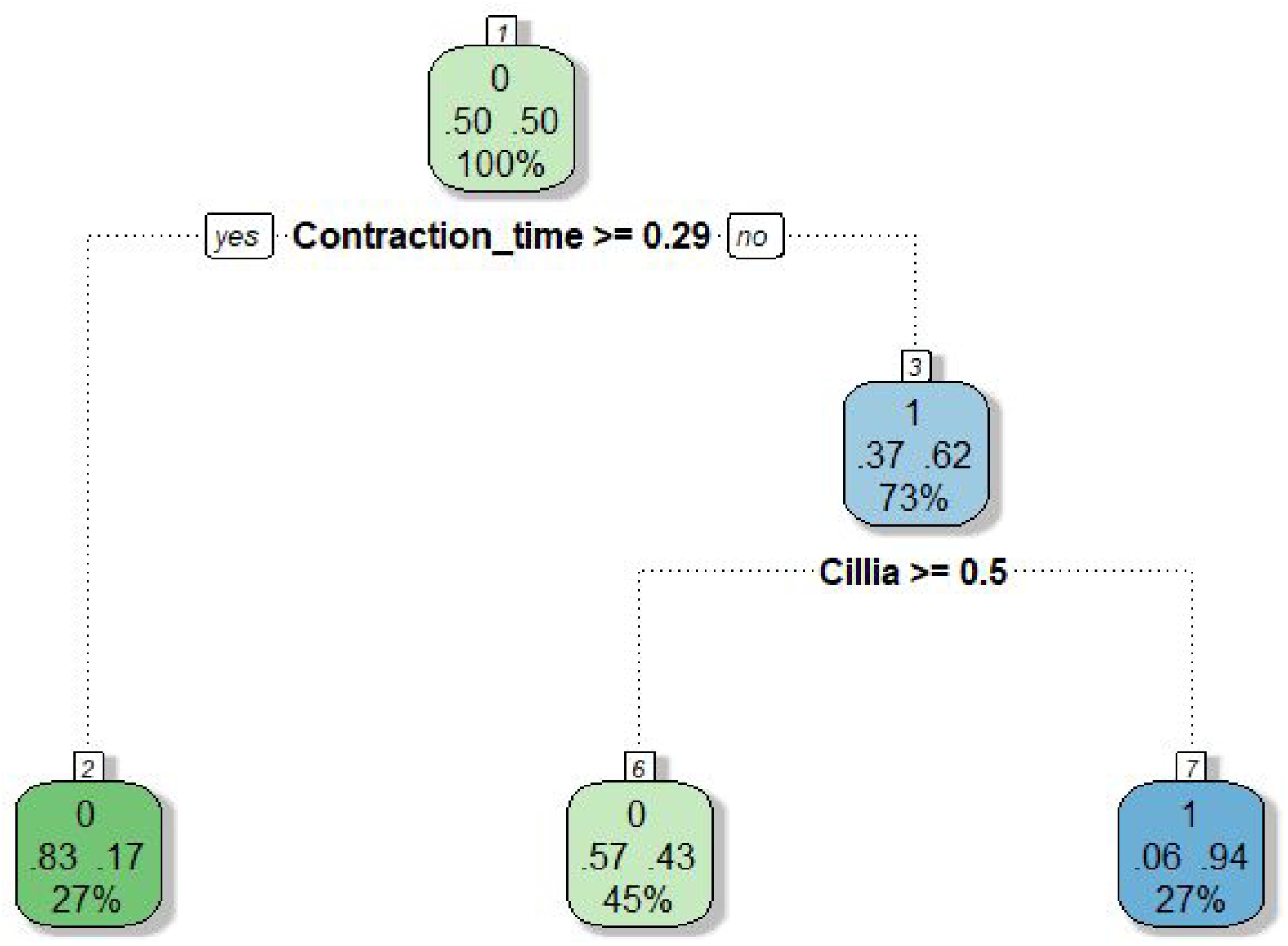
Tree representation of the classification process performed by Decision Tree model on training dataset. At each branch, a decision was made based on the feature printed to classify the dataset into class 0 (did not leave) or class 1 (did leave). The model used Contraction_time and then Cilia to classify the whole training dataset in 2 steps. Each box represent a group being classified, showing the class predicted, the probability of correct prediction, and size of the group as a percentage of the whole training dataset. (eg. in box 2, probability of an organism in this group belonging to class 0 is 0.83, and the group contains 27% of the whole training dataset.)

The second model we used was Random Forest. It is a different tree-based model, in which multiple trees are generated at the same time instead of the single tree approach that Decision Tree uses. Each tree will categorize a subset of the training dataset based on a random selection of input features, but only the variable with the highest association with the target will be chosen. The predictions generated from this collection of decision trees will be analyzed further and the class of highest “votes” will be chosen as an overall result. Random Forest model also allows variable importance to be assessed and extracted in the format of a ranking (Fig. 6). The mean decrease Gini measures the average reduction in purity of splitting events. Features that are highly correlated with the outcome seem to contribute more to the variation, hence, usually found most useful for prediction as they tend to help splitting mixed nodes into those with higher purity (indicated by Gini index). Here, the top three variables chosen by the model are, again, time spent in contraction, cilia reversal, and number of contractions, in order of decreasing importance.

**Fig 6.**
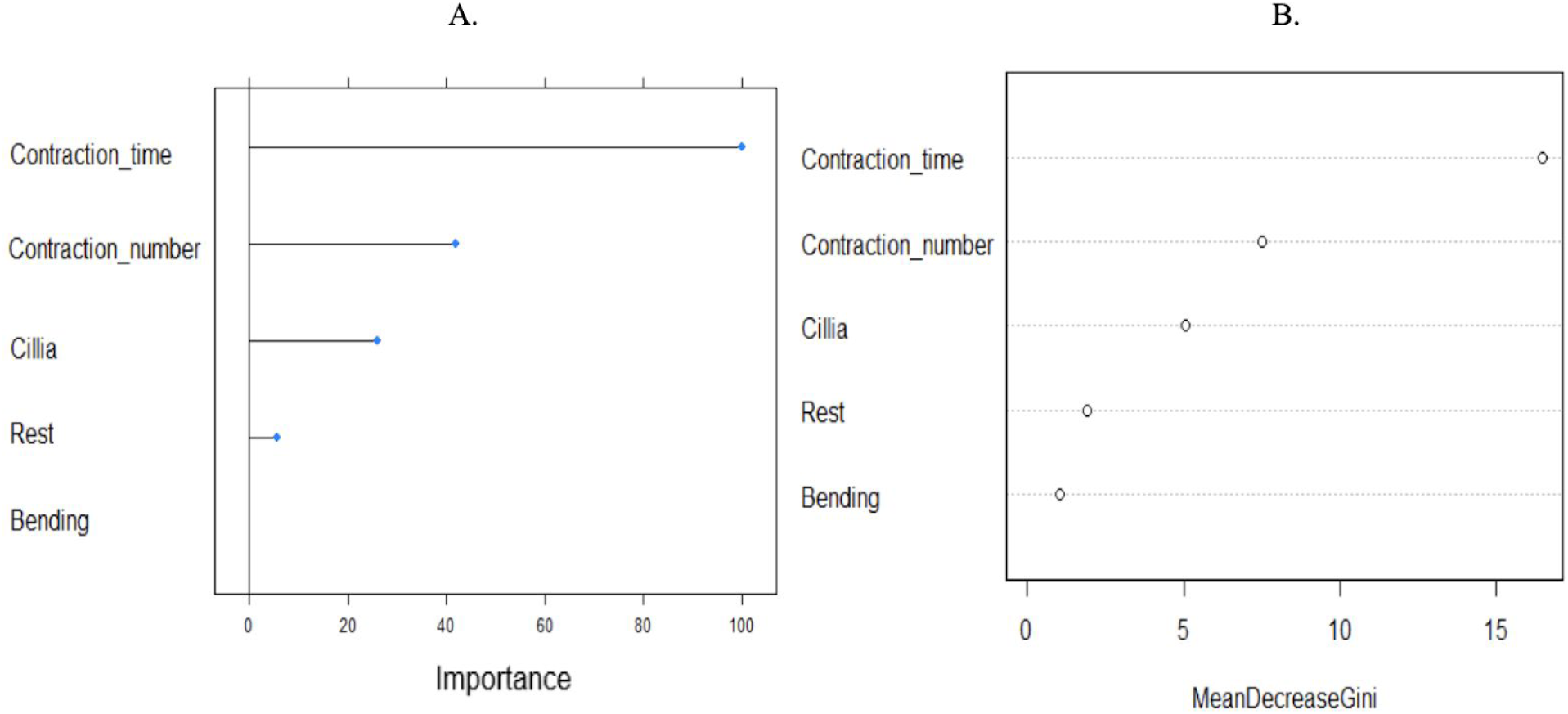
Variable Importance extracted from Random Forest model. A. The degree of importance of each input feature in classifying the training dataset, represented as percentage: Contraction_time being the most important variable with 100% importance; followed by Contraction_number, Cilia, Rest and Bending B. Contribution of each feature to the mean decrease in Gini index (indication of node impurity). The higher the decrease in Gini index, the higher the contribution of the feature to the nodes homogeneity.

The algorithms performed by decision tree are relatively clear, as we can examine the computations generated by the model, whilst random forest is more complicated with a big forest of deep trees. To gain a full understanding of the decision process by examining each tree is almost impossible. Although these approaches are easy to interpret and provide straightforward visualizations, their level of depth in inferring relationships and patterns from the data set is relatively poor. Since they mainly pick out variables that have the most significant impact on making predictions, they capitulate to capture other finer and more subtle details from the training data set involving the rest of the input variables. It does not make sense to conclude that the first two reactions in the series of five are insignificant or irrelevant as the organisms would not have done so otherwise.

Hence, feed-forward neural networks were used to further investigate the series of response in *S. roeselii*. These networks are made up of structured layers of computational neurons called ‘perceptrons’ (Fig. 7), mimicking actual biological input and activation architecture of real neurons. An Input layer takes in information from all training features, which is then passed on to hidden layer(s). There can be more than one hidden layer, and the perceptrons architecture can be customized. The more hidden layers there are, with highly intricate connecting algorithms, the more complex the network is, and hence the “deep” learning. These layers then perform computations that cannot, yet, be understood. The output of one layer is used as an input for the next layer. Eventually it will reach the final output layer, where the predictions are made.

**Fig 7.**
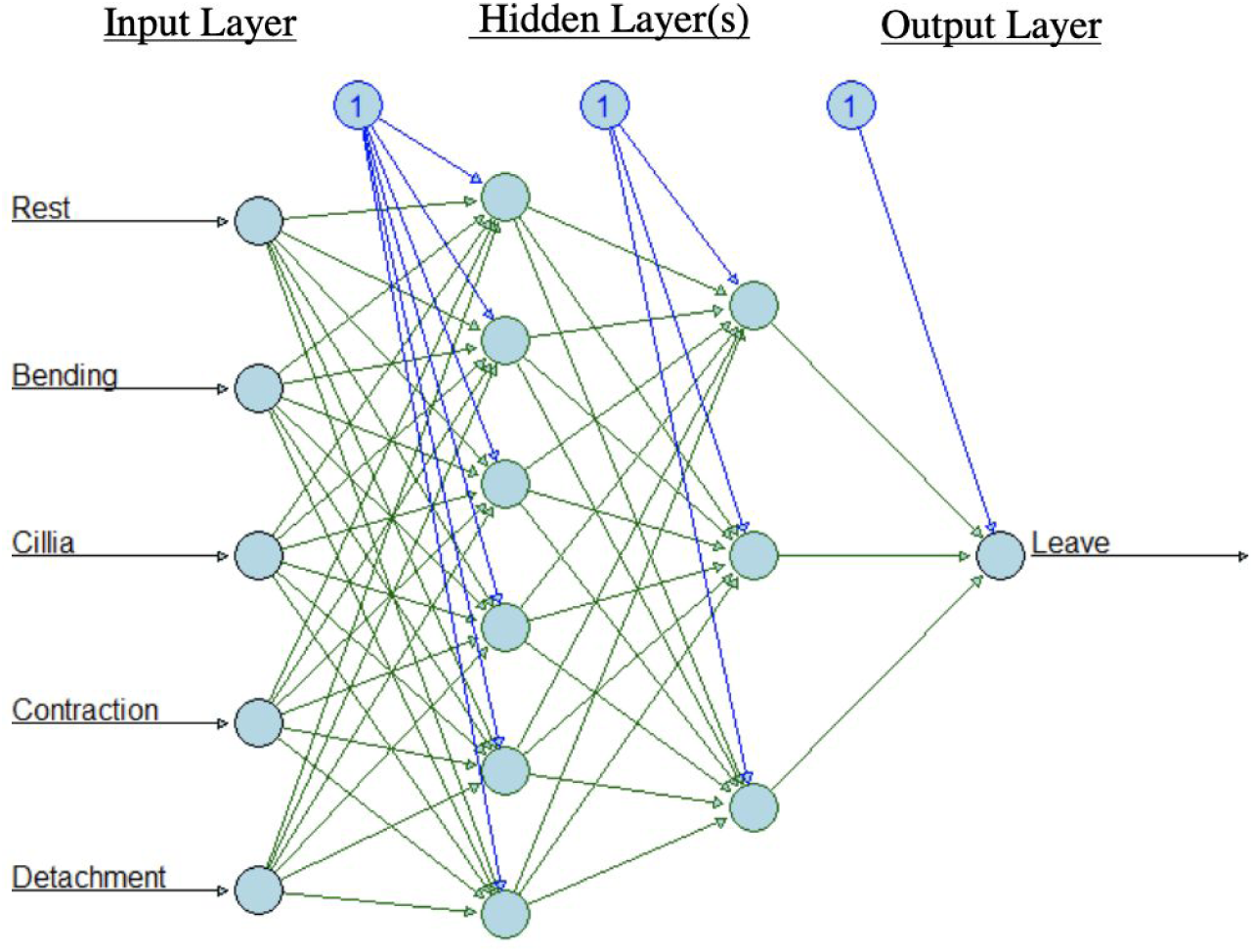
Schematic representation of an Artificial Neural Network (ANN) architecture. This particular ANN has an input layer with 5 perceptrons (blue circles) for 5 input features; 2 hidden layers: the first one with 6 perceptrons, and the second one with 3 perceptrons; and 1 output layer with 1 perceptron for binary classification outcome

Three feed-forward models were compiled with different hidden layer architecture and complexity (Table 1). After being trained, the performance of these models on a novel dataset can be evaluated using several different metrics. Some of these are summarized in Table 1, as well as being demonstrated through the Receiver Operator Curves (ROC) curves (Fig. 8). Accuracy implies the ratio of number of correct predictions out of all predictions made, and therefore, it seems like the higher the accuracy, the better the performance intuitively. Yet, this is not always the case, especially if statistical tests show that it is not significant, or if the dataset is imbalanced. F1 scores are one of the popular metrics used for evaluation of machine learning algorithms. It is the measure of the model’s precision and robustness. Generally, the higher the F1 score, the better the performance as it shows that not only could the model make predictions with adequate accuracy, but it also did not miss out too many difficult instances. Other evaluation metrics include the model’s sensitivity (ie. True Positive Rate or TPR) and specificity (can be interpreted by False Positive Rate or FPR), which can be summarized into Receiver Operating Characteristic – ROC curves (Fig. 8). TPR tells the proportion of class 1 samples that were correctly classified, whereas FPR tells the proportion of samples classified as class 0 that are False Positives. When the ROC curve is above the diagonal line, it means that the proportion of correctly classified samples in class 1 is greater than the proportion of samples that were incorrectly classified as class 0. The area under this curve (AUC) is particularly widely used as an evaluation metric for binary classification problems, which is very applicable in our experiment. AUC helps to compare different ROC curves for multiple machine learning models, and therefore, provides a measurement of model performance. Typically, the higher the AUC value, the better the performance.

**Table 1.** Metrics used to evaluate performance of 3 multilayer neural network models

|  | **Model 1** | **Model 2** | **Model 3** |
| --- | --- | --- | --- |
| **Hidden Layer Architecture** | 1 hidden layer<br>6 nodes | 3 hidden layers<br>20, 10, 6 nodes<br>respectively | 2 hidden layers<br>9, 18 nodes<br>respectively |
| **Accuracy ( %)** | 58.93 | 64.29 | 58.93 |
| **F1 Score** | 0.4091 | 0.28571 | 0.4889 |
| **Specificity** | 0.5952 | 0.76190 | 0.5238 |
| **Sensitivity** | 0.5714 | 0.28571 | 0.7857 |
| **AUC ( %)** | 63.2 | 66.2 | 73.0 |

**Fig 8.**
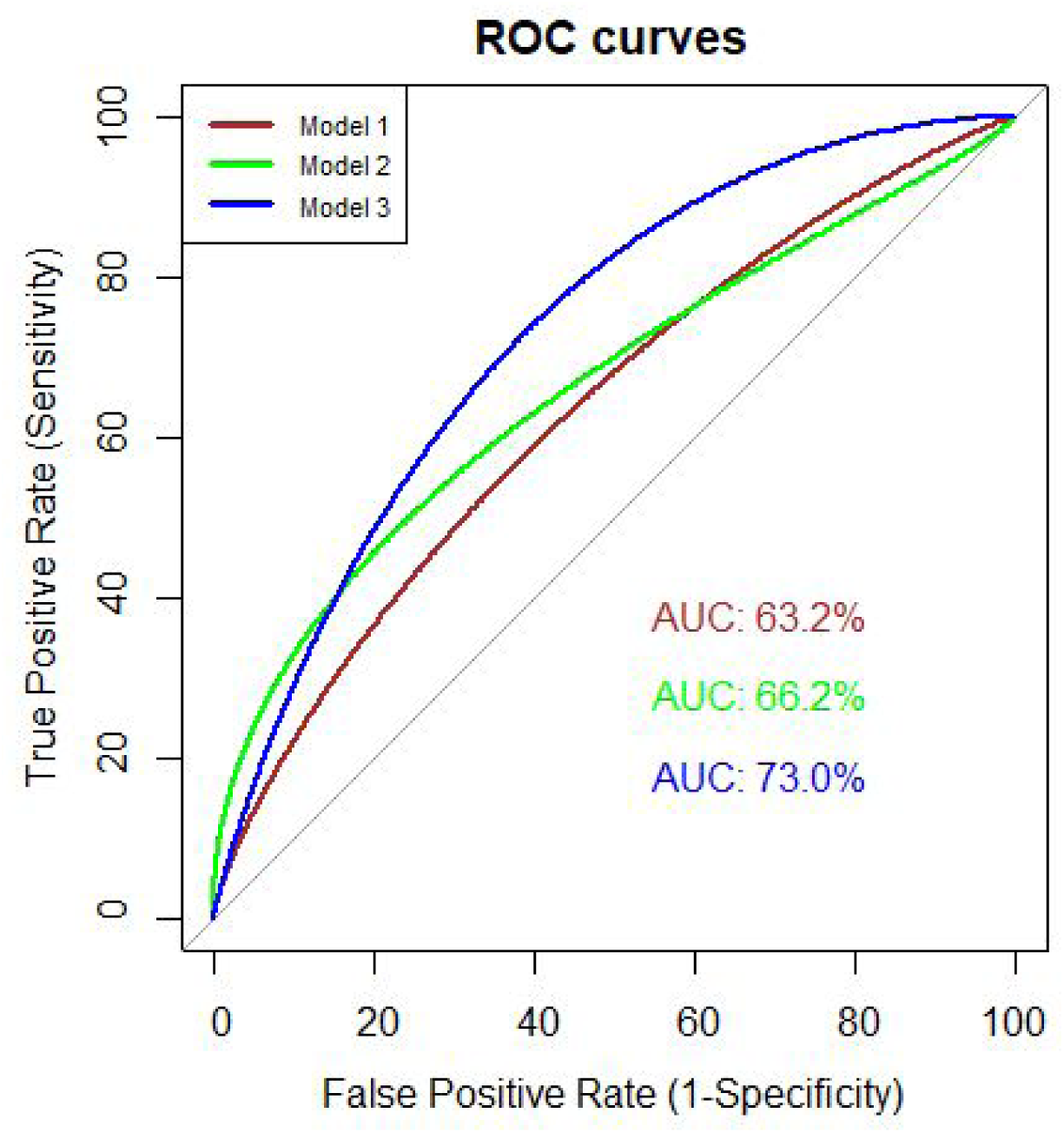
Receiver operating characteristic (ROC) curves (smoothened) for the three multilayer neural network models. These curves were generated and the corresponding area under the curves (AUCs) were calculated using R Studio, showing that model three gives the best performance overall at classifying the dataset.

Taking all these metrics into account, we have concluded that model 3, with the average level of network architecture complexity (of the 3), was the best model used for inferring meaningful patterns from the *S. roeselii*’s behavioural dataset to make predictions. It is important to emphasize that a simple ANN like model 1 is of no benefit. Even the best model can only produce roughly 59% accuracy. Yet, these ANNs contain many more “neurons” than the organism – *S. roeselii* is a single-cell aneural ciliate. What does this really mean? How can we can unravel the mechanistic details and computations that a highly complex brain does when we are not yet able to fully understand what a simple organism like *Stentor* is doing?

These results have highlighted the high level of complexity in the behaviours of *Stentor roeselii* in response to external stimulation. It cannot be fully explained by habituation, sensitisation, nor operant behaviour. Our machine-learning based models, though impressive in modelling activities of neural systems like the brain (Savelli & Knierim 2018; Yang et al. 2019), have been largely unsuccessful when applied here.

These aneural organisms exhibit fascinating behaviours that are considered learning, yet do not possess the complex neural networks of higher animals. Their ability to learn from environmental factors and respond appropriately would suggest the requirement for a different, but functionally equivalent network. Bray has argued extensively in his book “Wetware” (Bray 2009) how a system of protein molecules can perform all the tasks needed for a cell to sense and respond to its environment. The switching in behaviour of *S.roeselii* indicates an adaptational change in its internal state – i.e. the state of existing internal molecular networks (since the time scale is too short to allow for modifications in gene expression). The organism is changed by its previous experience, implying some form of memory. Molecular networks in single-cell organism and neural circuits in higher animals may have been independent evolutionary events but may also be fundamentally related.

And here comes the immense power of studying underlying mechanisms in lower single-cell organisms. It might seem counter-intuitive initially, but the pathway regulating E. coli’s chemotaxis behaviour – possibly the most well characterized pathway in biology to date – has taught us many critical lessons that can be applied to higher organisms, including humans, which also helped to develop general biological principles. As mentioned previously, *Stentor* is not the only example of a unicellular organism displaying learning behaviours but many more aneural organisms (Applewhite 1979; Applewhite & Gardner 1973; Reid et al. 2012; Shirakawa et al. 2011; Eisenstein 1975) have been extensively studied, leading to surprising results. Exploring the mechanistic details underpinning these behaviours can reveal profound insights into how neural circuits function in higher animals.

These ANNs, however, can be further developed, both by improving the experimental design and establishing more fine-tuned, advanced models. Our training dataset was relatively small and highly imbalanced. We addressed this issue by applying down-sampling method, however, it resulted in an even smaller set. It would be much more beneficial to have a bigger data set of higher control and filming quality, as these directly impact the performance of the ANNs. Equally, these ANNs can be further developed to a higher level of complexity, with better suited and tuned parameters to increase their performance. This would allow more features to be extracted from the raw data. The best source to search for guidance to improve computer-based models is from our understanding and knowledge of the underlying biology (Dasgupta et al. 2017).

We now know that learning and memories are required even at single-cell level. All living cells must have an awareness of their immediate environment to a certain extent. Components of the immune system, like macrophages or neutrophils, are constantly required to learn and form memories (Prentice-Mott et al. 2013). We are now in a much better position to study these intriguing behaviours in aneural organisms, or single-cell behaviours generally, which were undreamt of by scientists like Jennings a hundred years ago. With the abundance of advanced biochemical and genetic tools, we are able to unravel the molecular circuits inside these single cells in much more detail. These results are then combined with computational simulation and computer-based artificial intelligence, creating a powerful synergistic effect which will one day decode the mystery behind these observations. One might question the validity of using ANNs in modelling biological molecular networks. Despite many differences in details, general principles and properties are still shared (Bray 2009). This research contributes significantly to our ultimate goal of elucidating general principles of computations taking place in the brain (Carandini 2012). Nonetheless, the biggest drawback of this approach is, indeed, its black-box nature. There have been multiple efforts recently to open this black box and reveal the computations performed by ANNs (Castelvecchi 2016; Zhang et al. 2018). As Richard Feyman’s famously said – whether it be ANN computations or living cellular behaviours – “What I cannot create, I do not understand”.

## Acknowledgments

We thank Dr. M. Oliva, Department of Genetics, University of Cambridge for help with the set up of experimental apparatus.

### Funding

KMT is funded by Cambridge Trust Scholarship and Trinity Overseas Bursaries; SP is funded by the Cambridge-DBT lectureship.

## Author contributions

KMT performed all the experiments, analyzed the data and extensively contributed to the draft. MTW helped with the microscopy experiments and ordering Stenors. He also critically evaluated the analysis, results and manuscript. SP designed and supervised the project, analysed the data, and wrote the manuscript.

## Data and materials availability

All videos and raw data will be made available.

## Material Methods

### S.roeselii source and maintenance

Cultures of *S.roeselii* were purchased from Sciento (Manchester, UK) approximately every week over a period of 1 month. Sciento harvested the organisms from a pond on the property of Whitefield Golf Club (83 Higher Lane, Whitefield, Manchester, UK). In the lab, *S. roeselii* was maintained in well-aerated glass flasks in pond water, which were kept mainly in the dark, with partial sunlight. All behavior experiments were performed on organisms purchased no more than seven days beforehand.

### Micro-stimulation apparatus and set-up

Custom-built apparatus to deliver controlled pulses of polystyrene beads directly near the mouth of the organism was used. A small clamp was placed next to the microscope. The microinjection glass needle was loaded with the suspension of fluorescent red latex beads (Fluorescent-red, carboxylate-modified polystyrene beads in aqueous suspension with 0.1% NaN3 (Sigma-Aldrich; mean diameter 2 μm)) and connected to an elevated reservoir of distill water. The needle was then held next to the microscope by the clamp.

One drop of *S. roeselii* culture was placed on a glass slide for each observation. The droplet culture was allowed to settle down for few minutes. The microinjection needle was positioned next to the mouth of the organism by hand, and its position was adjusted as needed throughout the experiment using the clamp. Short pulses of beads were generated as a gravity-flow with the opening and closing of a two-way stopcock connected to the bottom of the reservoir, or adjusting the height of the reservoir.

### Microscopy

All images were acquired using a Leica MZ16F Stereoscope equipped with a 11.25x objective lens and a QImaging Retiga 2000R monochrome camera. Images were collected at a rate of 15 frames per second for time lapse experiments, using an exposure time of 16.184 ms using Micro-Manager (Edelstein et al., 2014). All microscopy experiments were performed at the Imaging Facility, Zoology Department, University of Cambridge.

### Video analysis

29 collected videos were analyzed and investigation was performed on 188 individual organisms. Detailed description of *S. roeselii’*s behaviours were recorded along with the corresponding time in the video. A video would be terminated if: (1) all sessile *S. roeselii* “decided” to leave and swam away, or (2) the specimen dried out even though there were still organisms being attached to the piece of algae.

### Modelling

#### All modelling was done using R Studio

Raw data collected from video analysis were converted into time spent (1) at rest, (2) bending, (3) reversing cilia, (4) contraction, (5) number of contractions, and (6) leave (or did not leave). These data were then scaled and randomly split into a training dataset (70%) and a testing dataset (30%). A down-sampling method was applied to the training dataset to ensure a 1:1 ratio between 2 classes (“Leave” and “Did not leave”). New training dataset is then shuffled before being used to train the models.

Decision Tree and Random Forest models were compiled using rpart, randomforest (respectively), and caret packages in R. 10-fold cross validation with 10 repetitions were used to train the models.

Feed-forward neural networks were compiled, trained, and evaluated using keras package.

---

**Fig S1.**
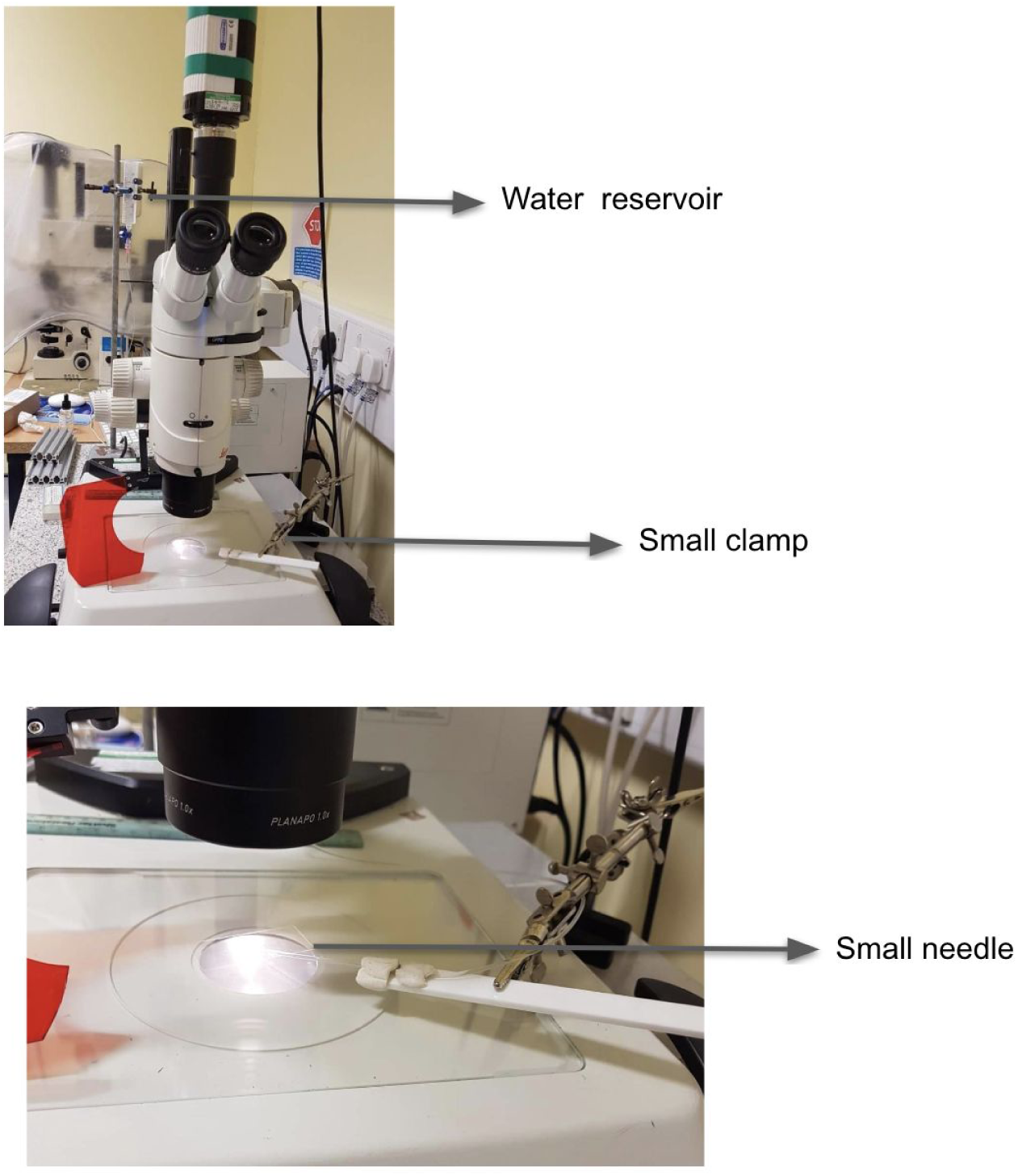
**A. Micro-stimulation apparatus and set-up.** Picture of the whole apparatus set up **B. Glass needle set-up.** Picture of the zoomed in section where showing the glass needle.

